# Stimulus blanking reveals transsaccadic feature transfer

**DOI:** 10.1101/819110

**Authors:** Lukasz Grzeczkowski, Heiner Deubel, Martin Szinte

## Abstract

Across saccadic eye movements, the visual system receives two successive static images corresponding to the pre- and the postsaccadic projections of the visual field on the retina. The existence of a mechanism integrating the content of these images is today still a matter of debate. Here, we studied the transfer of a visual feature across saccades using a blanking paradigm. Participants moved their eyes to a peripheral grating and discriminated a change in its orientation occurring during the eye movement. The grating was either constantly on the screen or briefly blanked during and after the saccade. Moreover, it either was of the same luminance as the background (i.e., isoluminant) or anisoluminant with respect to it. We found that for anisoluminant grating, the orientation discrimination across saccade was improved when a blank followed the onset of the eye movement. Such effect was however abolished with isoluminant grating. Additionally, performance was also improved when an anisoluminant grating presented before the saccade was followed by an isoluminant one. These results demonstrate that a detailed representation of the presaccadic image was transferred across saccades allowing participants to perform better on the trans-saccadic orientation task. While such a transfer of visual orientation across saccade is masked in real-life anisoluminant conditions, the use of a blank and of isoluminant postsaccadic grating allowed here to reveal its existence.

**Significance statement:** Static objects are perceived as not moving across eye movements despite their visual projection shifts on our retina. To compensate for such shifts and create a continuous perception of space, our brain may keep track of objects’ visual features across our movements. We found that shortly blanking a contrast-defined object during and after saccades allows to recover a detailed representation of its orientation. We propose that the transfer of visual content across saccades revealed with the use of a simple blank plays an important role in ensuring our continuous and stable perception of the world.

## Introduction

The mosaic of photoreceptors of the human retina is highly inhomogeneous (1). This results in high resolution vision in the center of the visual field and low resolution vision in its periphery (2). To account for that inhomogeneity, humans sample their environment by making frequent saccadic eye movements. But to make a saccade to an object of interest, the visual system must first select that object in between others. This selection is necessarily based on the low-resolution information available in the visual periphery that is replaced by the high resolution, foveal image at the end of the saccade (3). Thus, across saccades, the visual system receives two images of an object, a presaccadic low-resolution image in its periphery, and a postsaccadic high-resolution image in its fovea. The mechanism used by the visual system to create our perception of space constancy across these resolution changes is still today a matter of debate (4). In particular, an open question resides on whether the visual content of the presaccadic image can be transferred across saccades and to which extent it is integrated with the postsaccadic image (5, 6).

Most of the early studies on that question concluded that trans-saccadic perception relies on visual short term memory with a transfer of abstract, relational and structural aspects of visual stimuli across saccades (7). Moreover, it was proposed that the detailed presaccadic information, the stimulus features and its precise position are not transferred, nor integrated across saccades (8–11).

Recent studies however changed this view, showing that low level, pre- and postsaccadic image visual features such as orientation (12–14) or color (15–17) are indeed partially transferred and integrated across the saccade, probably using a presaccadic remapping mechanism (18–22).

Importantly for the context of the present study, it was also shown that relatively precise information of object location can be maintained across saccades and be almost perfectly recovered if the attended object is briefly blanked at the onset of the saccade (23, 24). Interestingly, this blanking effect turned out to be strongly reduced for low contrast and isoluminant stimuli (25), suggesting that the mechanism used to transfer position across saccades relies on luminance contrast information. In line with the above findings, Paeye, Collins, & Cavanagh (2017) showed that two sets of lines presented before and after eye movements could be perceptually fused to form a letter. They interpreted their results as evidence of a spatiotopic visual persistence generated by neural remapping processes (26).

Here, we hypothesize that the blanking effect (23) relies on the transfer across saccades of a remapped, representation of the presaccadic image. To test this hypothesis, we evaluated participants’ ability to judge a change of a grating orientation occurring during a saccade (Figure 1A-B). We predicted that the addition of a short blank during and after the saccade would help participants to judge the trans-saccadic orientation change. We hypothesized that with a blank, participants could base their judgment on apparent motion (27) occurring between the transferred presaccadic orientation representation and the postsaccadic one, directly seen on the screen. Moreover, given that the blanking effect for object location depends on the luminance contrast of the stimulus (25), we predicted that only judgments made for anisoluminant stimuli would benefit from the blank. Our results confirmed our hypotheses, with strong improvement of discrimination thresholds when a blank was introduced at the saccade onset. Moreover, detailed recovery of the presaccadic object orientation across saccades was observed only for anisoluminant, but not for isoluminant stimuli. Remarkably, in the No-blank condition, performance was also improved when presaccadic anisoluminant gratings were followed by isoluminant postsaccadic gratings.

**Figure 1.**
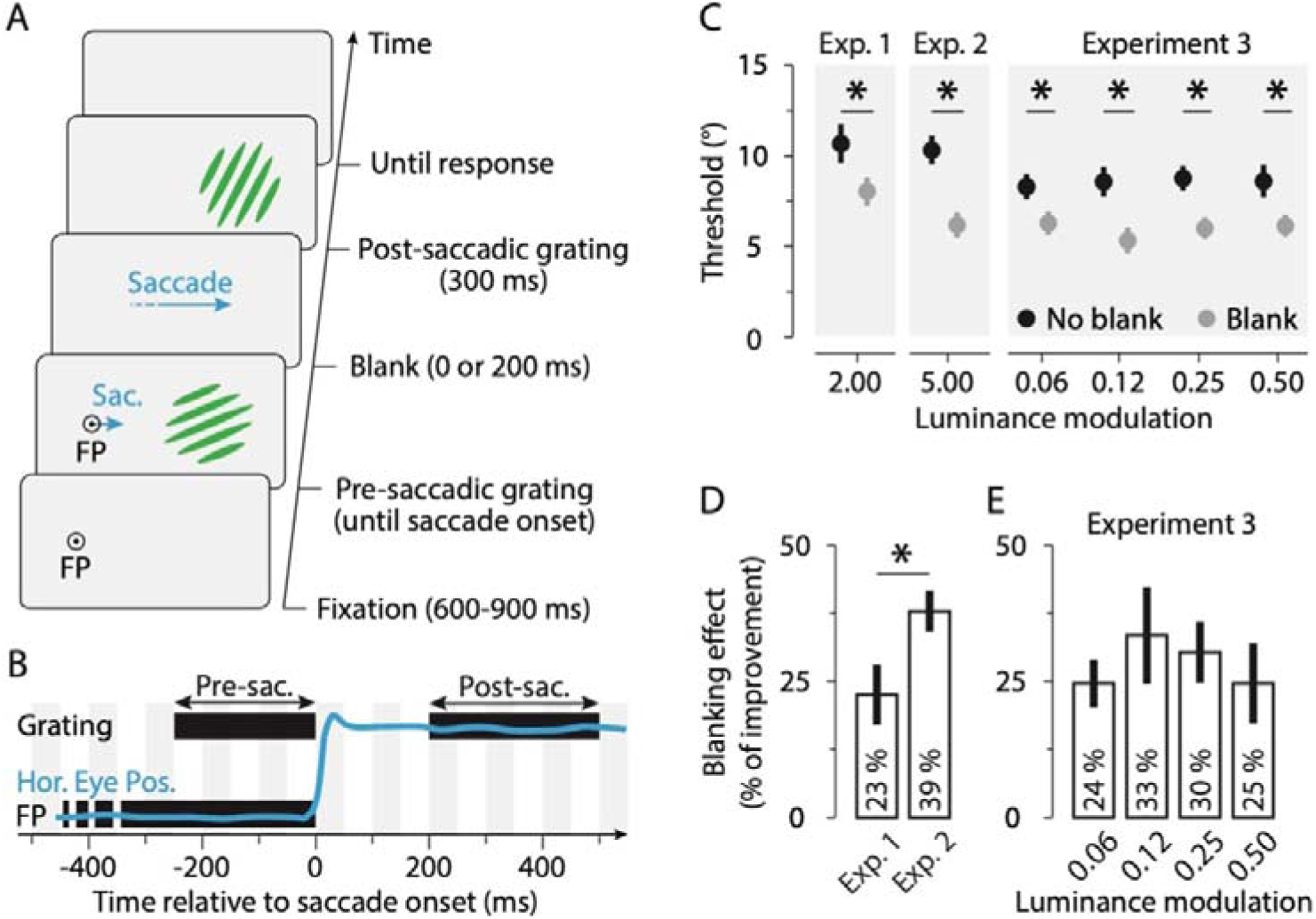
Procedure and orientation blanking effect. **A**. In all experiments, after fixating the fixation point (FP), participants made a saccade towards a peripheral presaccadic oriented grating. The saccade triggered the presentation of the postsaccadic grating, rotated clockwise or counterclockwise relative to the presaccadic one. The postsaccadic grating was presented either immediately (No Blank condition) or 200 ms after a period during which the screen was blanked (Blank condition). Participants reported the direction of the orientation change at the end of each trial. **B**. Time course of a trial. The orange line shows horizontal eye position (Hor. Eye Pos.) relative to saccade onset in a trial in which a blank was used. Black rectangles illustrate the timing of the pre- and postsaccadic gratings and the fixation point (FP). **C**. Average orientation threshold angle observed with anisoluminant gratings in Exp.1, Exp. 2 and Exp. 3, in the No blank (black) and Blank (gray) conditions, respectively. **D-E**. Bars show the blanking effect (see methods) observed in the Exp.1, Exp.2 and Exp. 3. Error bars show SEM and asterisks and lines show significant comparisons (p < 0.05).

Together, these results suggest the existence of a mechanism allowing the transfer and the integration of visual features across saccades. We here made such a mechanism directly apparent by the use of a short blank during and after the eye movement.

## Results

We first aimed at determining how large the change of the grating orientation must be such that participants can correctly discriminate its orientation change across saccades. We found that with anisoluminant gratings, that is gratings with a twofold (Exp. 1) or fivefold (Exp.2) luminance relative to the background, participants needed a rotation change of about 10.5° relative to presaccadic grating angle to discriminate the change correctly in 75% of the trials (Figure 1C, black; Exp. 1: 10.68 ± 1.17, Exp.2: 10.29 ± 0.86).

We found that orientation threshold could be strongly reduced if a 200 ms blank followed the saccade onset. When a blank was used, participants needed only a rotation of ~7° to perform the task (Figure 1C, gray; Exp. 1: 8.05 ± 0.85, *p* = 0.0062, *d* = 0.81; Exp. 2: 6.17 ± 0.52, *p* < 0.0001, *d* = 1.84). On average, orientation discrimination thresholds were improved by about 30% with the use of a blank, an improvement slightly less pronounced with twofold (Figure 1C, Exp. 1: 23.20 ± 5.73%) than with fivefold luminance gratings (Figure 1C, Exp. 2: 38.80 ± 3.91%, *p* = 0.0048, *d* = 0.92). Interestingly, as a blank is by definition an absence of visual stimulation, these first results suggest that the transfer of the presaccadic orientation is partially masked by the presentation of the postsaccadic grating without a blank. In other words, the blank allows to better recover the feature content of the grating orientation across saccades by unmasking the presaccadic transferred information.

We tested whether the blanking effect occurs for much less luminanant gratings, i.e., gratings having 6, 12, 25 and 50% more luminance than the background (Exp. 3). We reproduced the orientation blanking effect at each stimulus luminance condition, with significant improvement of the orientation threshold within the same range as the Exp. 1 and Exp. 2 (Figure 1D-E, 0.0008 > *ps* > 0.0001, 1.63 > *ds* > 0.86). These results suggest that the orientation blanking effect is a robust phenomenon which reaches a plateau already at 6% of luminance.

While we systematically found an orientation blanking effect with anisoluminant gratings, we also tested and analyzed three other conditions in which we varied the luminance of the pre- and postsaccadic grating (Figure 2A). First, when both the pre- and postsaccadic gratings were isoluminant, i.e., gratings only defined by a change in color relative to the background, there was no difference in orientation threshold when comparing trials with (Figure 2B: pre-sac iso & post-sac iso, Exp. 1: 9.01 ± 0.83; Exp. 2: 7.96 ± 0.97) and without blank (Figure 2B: pre-sac iso & post-sac iso, Exp. 1: 9.56 ± 0.89, *p* = 0.42, d = 0.20; Exp. 2: 8.13 ± 0.77, *p* = 0.45, *d* = 0.06). Importantly, these results show that unmasking the transferred targets representation by blanking has no effect. This suggests that the precise representation of the presaccadic grating is only transferred across the saccade if the presaccadic image is defined by luminance contrast (25).

**Figure 2.**
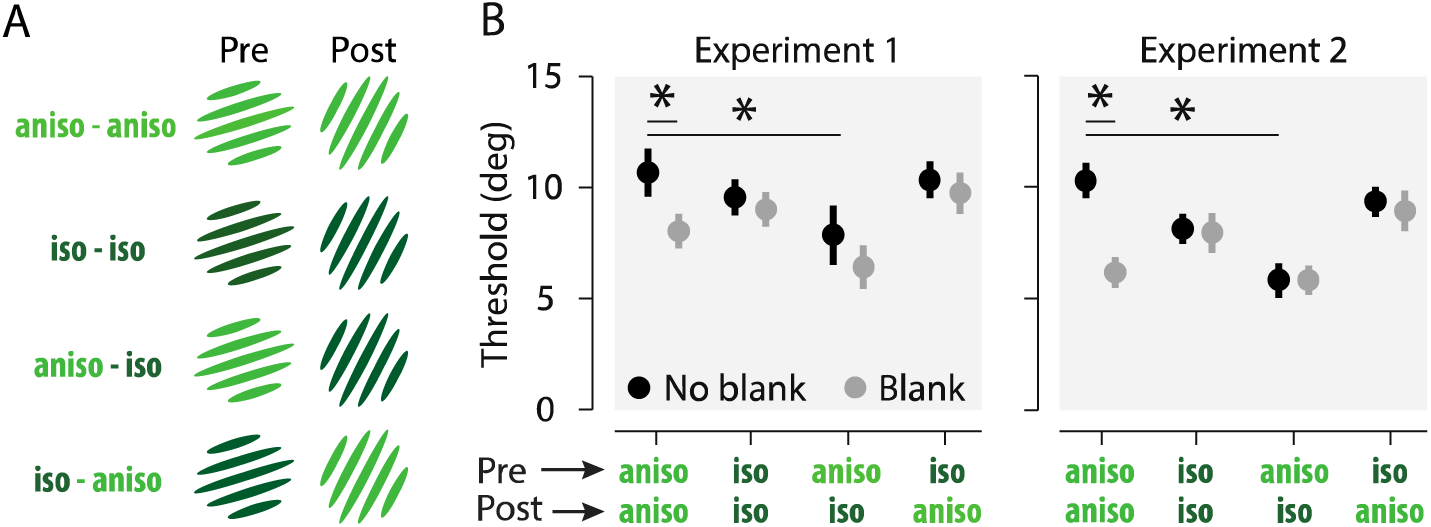
Effect of luminance on the orientation blanking effect. **A**. Combinations of pre- and postsaccadic gratings used in Exp. 1 and Exp. 2. **B**. Mean thresholds observed as a function of the pre- and postsaccadic grating luminance, in the no blank (black) and blank (gray) conditions, respectively. Note that results with pre- and postsaccadic anisoluminant gratings are here replotted from Figure 1. Conventions are as in Figure 1.

Second, no blanking benefits were found in trials in which a presaccadic isoluminant grating was followed by a postsaccadic anisoluminant one. In this condition, there was no improvement of orientation discrimination threshold when comparing trials with (Figure 2B: pre-sac iso & post-sac aniso, Exp. 1: 9.75 ± 1.00; Exp. 2: 8.93 ± 1.00) and without a blank (Figure 2B: pre-sac iso & post-sac aniso, Exp. 1: 10.91 ± 0.91, *p* = 0.26, *d* = 0.30; Exp. 2: 9.36 ± 0.63, *p* = 0.38, *d* = 0.21). This result further demonstrates that, similarly to the previous condition where pre- and postsaccadic gratings were both isoluminant, the transsaccadic transfer of orientation depends on the luminance contrast of the presaccadic image.

Finally, we tested a condition in which a presaccadic anisoluminant grating was followed by a postsaccadic isoluminant one. Here again, we didn’t find a blanking effect when comparing trials with (Figure 2B: pre-sac aniso & post-sac iso, Exp. 1: 6.44 ± 1.10; Exp. 2: 5.83 ± 0.61) and without a blank (Figure 2B: pre-sac aniso & post-sac iso, Exp. 1: threshold = 7.87 ± 1.46, *p* = 0.41, d = 0.35; Exp. 2: threshold = 5.80 ± 0.55, *p* = 0.87, d = 0.02). One could first conclude that also in this condition, a precise stimulus representation was not available to the participants after their saccade. However, orientation thresholds in this condition (pre-sac aniso & post-sac iso) were systematically lower than those observed with anisoluminant pre- and postsaccadic gratings in both experiments (Figure 2B; Exp. 1: *p* = 0.0004, d = 0.67; Exp. 2: *p* = 0.0001, d = 1.98). This suggests that a precise representation of the presaccadic grating is present after the saccade when an object defined by luminance contrast (but not color) is presented as a saccade goal. However, this transsaccadic image is normally masked by the content of the postsaccadic image, at least when this second image contains luminance contrast.

How precise is the transferred representation? To answer this question, we analyzed participants’ just noticeable differences (JND) and biases in discriminating the orientation change (see Methods and Figure 3). We found large JND improvements when both, the pre- and postsaccadic gratings were anisoluminant as compared to other conditions. This effect is made evident for a representative participant (Figure 3A, top-left panel), showing a steeper psychometric function in the Blank (gray symbols) than in the No blank (black symbols) condition. This led to significant reductions of the observed JNDs when both, the pre- and the postsaccadic gratings were anisoluminant, for the No blank (Figure 3B; Exp. 1: JND = 16.25 ± 1.95; Exp. 2: 12.96 ± 1.34) and Blank conditions (Exp. 1: JND = 10.96 ± 0.99, *p* < 0.0002; Exp. 2: 8.15 ± 0.63, *p* < 0.0001). Interestingly, such improvement of discrimination performance was made apparent by the use of blank without change of participant judgment bias when comparing the No blank (Figure 3C; Exp. 1: Bias = 0.31 ± 0.07; Exp. 2: Bias = 0.50 ± 0.10) and the Blank condition (Exp. 1: Bias = 0.57 ± 0.08, *p* = 0.39; Exp. 2: Bias = 0.40 ± 0.08, p = 0.29). These results contrasted with what observed within our three other luminance conditions, for which we found that both the participants’ JNDs (Figure 3B; Exp. 1: 0.87 > *ps* > 0.08; Exp. 2: 0.93 > *ps* > 0.49) and biases (Figure 3C; Exp. 1, 0.73 > *ps* > 0.07; Exp. 2, 0.82 > *ps* > 0.14) stayed constant when comparing the No blank and the Blank conditions. Together these results suggest that the transferred representation is more precise than indicated by the normal, No-blank condition. When available, it rendered the orientation discrimination task easier over the whole range of the tested orientation changes without affecting participant bias.

**Figure 3.**
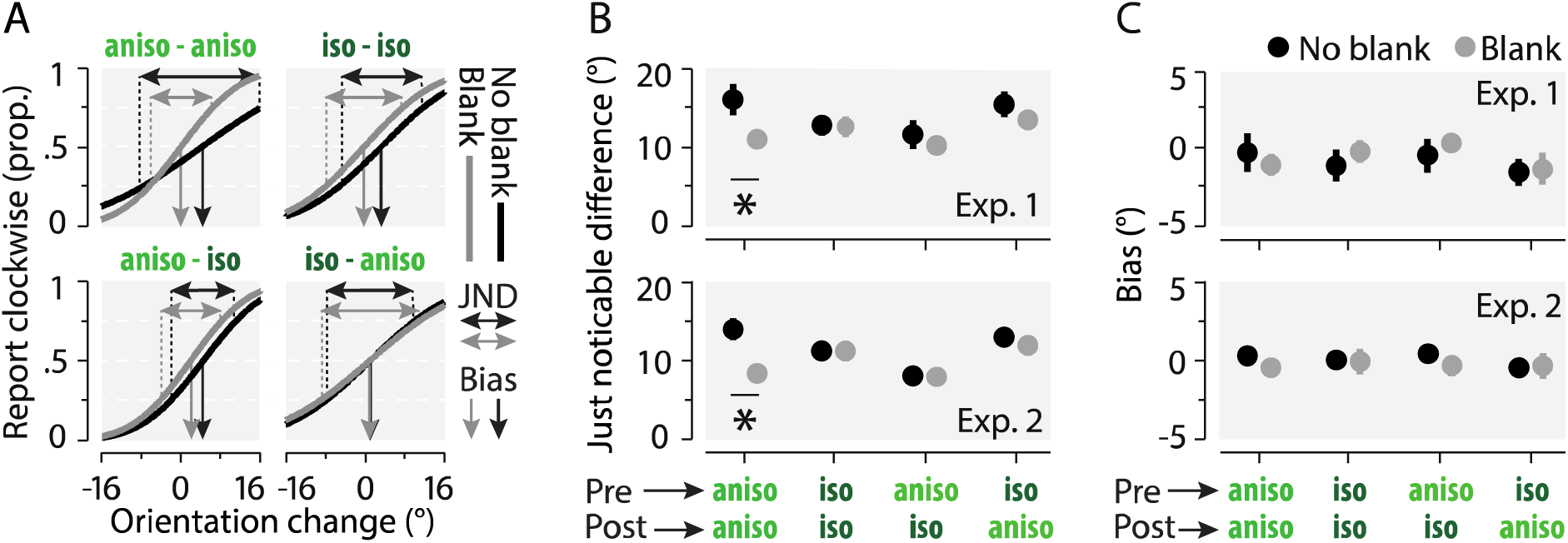
Just noticeable differences (JNDs) and biases as a function of the gratings’ luminance. **A.** Psychometric functions of a representative participant showing the proportion of clockwise responses as a function of the orientation change of the gratings across saccade in each luminance (see panel titles) and blanking condition (Blank and No blank in gray and black, respectively). Horizontal and vertical arrows show JNDs and biases, respectively for the blank (gray) and no blank (black) conditions. **B-C.** Average observed JNDs (B) and biases (C) across participants in Exp. 1. (upper panels) and Exp. 2 (lower panels). Conventions are as in Figure 1.

## Discussion

Early work on trans-saccadic perception suggested that space constancy is achieved with transsaccadic transfer of visual features leading to spatiotopic visual persistence across saccades (28). Interestingly, these studies reported that participants experienced seeing the stimuli after the saccade, even if they had been removed from the screen (28). As persistence could have been due to the screen used (29) this proposal was for long abandoned (8–10, 29, 30). Nevertheless, a new series of studies showed the existence of a partial but systematic transsaccadic integration of visual features across saccades (12–14, 16, 22, 31). Moreover, the proposal of a spatiotopic visible persistence was recently revisited in a study using subjective reports (26) and EEG decoding (32).

Using psychophysical methods, we measured the potential perceptual effects of a spatiotopic visual persistence across saccades. As reported earlier (12–14), we found that visual orientation information is partly transferred across saccades, allowing participants to report the change in orientation of a grating across eye movements. Importantly, we showed in three experiments that briefly blanking the screen during and after the saccade allowed participants to recover the orientation of the presaccadic grating, as made evident by the strong and systematic improvement of participants’ orientation thresholds and JNDs. Moreover, while we found the strongest orientation blanking effect with gratings of high luminance (Exps. 1 and 2), the effect seemed to reach its maximum with low luminance stimuli already (Exp. 3: between 6% and 12% more luminance then the background), suggesting the generality of the underlying mechanism. These results differ from a previous report (25) in which the blanking effect for position had a gradual effect as a function of the contrast level. Nevertheless, with orientation change discrimination, we similarly found an absence of a blanking effect for isoluminant stimuli (25). This effect is particularly interesting as it suggests that feature transfer across saccades relies on the presence of luminance contrast in the presaccadic scene. Moreover, it demonstrates that the presence of a blank after saccade onset allows the recovery of presaccadic information that is usually masked by the postsaccadic image. In line with that result, participants better reported the orientation change when the presaccadic anisoluminant grating was followed by an isoluminant one, irrespective of the presence of a blank. This suggests that the transferred orientation across saccades wasn’t masked as effectively by a postsaccadic isoluminant image as it was by an anisoluminant one. Finally, the changes in the just notifiable difference (JND) observed with anisoluminant gratings suggest that the transsaccadic judgment task was easier when a blank followed an anisoluminant image or when the postsaccadic image was isoluminant (Figure 3).

Our results suggest the existence of a visual mechanism allowing the transfer of visual content across the saccade. Indeed, our results suggest that the precise view of the presaccadic object, normally masked by the view of the postsaccadic object can be made directly visible to participants when a blank is used. We propose that this representation made visible by the blank induces a percept of apparent motion between the spatiotopic image of the presaccadic grating and the postsaccadic grating (27), making the transsaccadic discrimination task easier. Importantly, our results also imply that low-level visual features are transferred across saccades and that under normal conditions, the transsaccadic memory is, to a large extent, hampered by the postsaccadic scene. In this framework, blanking the postsaccadic scene postpones or avoids the masking of the transferred visual feature enabling access to its content maintained in memory (33).

It was recently shown that predictive coding can account for feature transfer across saccades (34, 35). Predictable objects were shown to be better detected (34) and to lead to decreased BOLD activity in the early visual cortex as compared to unpredictable ones (35). Predictive coding models assume that visual stimuli after being processed in high-level visual areas are fed back to low-level neurons in order to create visual predictions (36). In a similar manner, high-level processes can generate low-level percepts through visual imagery. Visual imagery can lead to the formation of a short-term sensory trace that can bias future perception (37) and even lead to visual perceptual learning (38). Besides visual imagery, ghostlike percepts such as observed in filling-in phenomenon, were shown to generate neural activity in the early visual cortex (39). We thus argue that, alike visual imagery and filling-in phenomenon, we here demonstrated a phantom-like featural representation that could result from predictive coding fed back to low-level visual neurons.

We here propose that the presaccadic stimulus generates first a feed forward propagation of the signal to higher-level visual areas, later fed back to early visual areas at predicted spatiotopic locations. This back-propagation occurs during the saccade and ends after the saccade offset. Accordingly, 50 ms after the saccade offset, the detection advantage of target predictability (34) and the blanking effect for target displacement begin to be effective (23). The predictive activation from the high-level areas would later excite low-level neurons which, similarly to visual imagery or filling-in phenomena would create the phantom-like, featural representation.

However, to be seen at the correct spatiotopic position in space despite change in retinal coordinates, featural predictions must be remapped between the visual neurons accounting for the different pre- and the postsaccadic retinal positions of the gratings. A transfer of neural information was demonstrated in higher level visual areas such as the FEF (40), parietal areas (20) and the superior colliculus (41) which feeds back visual position information to low-level visual areas involved in visual orientation processing (42). To allow a spatiotopic representation of visual features, remapping of orientation information must also take place. Up to date, such neural transfer of feature preference across saccades has not been found (43). Possibly, however, the feedforward signals of the postsaccadic image, akin our masking effect observed with postsaccadic anisoluminant gratings, could have hampered the measure of the feature transfer. Future work using our blanking paradigm may demonstrate the existence of feature specific, neural remapping mechanisms.

It was previously argued that only the position of attended objects would be transferred across the saccade (4, 5). However, our results as well as evidence of transsaccadic feature integration (12, 13) suggest that not only the position but also visual features are transferred. In this regard, others argued that head-centered, spatiotopic, or gaze-modulated mechanisms could explain space constancy and transsaccadic perception (6). While the visual system has been shown to be organized in retinotopic coordinates (44), our results suggest that at one stage of the visual hierarchy the featural content of the presaccadic image is maintained in space. Thus, our results support the existence of a neural mechanism allowing visual neurons to keep track of the orientation of the grating across saccades. However, today, there is no available electrophysiological evidence favoring this alternative.

Irrespective of the neural mechanism behind, our results argue in favor of a precise, phantom-like, featural representation potentially ensuring space constancy. However, our results also suggest that under natural, anisoluminant conditions, such representation is partially masked. While transsaccadic partial transfer of visual features might be sufficient for space constancy, it could also well be that feature information is generally not used across saccades or integrated with the postsaccadic information (12, 13). If it would be the case, maintaining the position of attended objects across saccade (4, 19) or assuming visual stability of the world could be sufficient to create an impression of stability. The transfer of visual features as observed here will then only reflect the fact that we experimentally broke this assumption using a blank. Future research taking advantage of the blanking paradigm will help to better understand transsaccadic mechanisms, for example by determining under which spatio-temporal conditions and for which stimulus attributes (e.g., color, size, spatial frequency) feature transfer can occur.

We here show that to compensate for retinal shifts across saccades and create our continuous perception of space, our brain may keep track of the visual features of objects across eye movements. We found that shortly blanking a contrast-defined object allows to recover a detailed representation of its orientation. We propose that the blank reveals a phantom-like featural representation allowing participants to accurately judge an orientation change across saccades.

## Material and methods

### Participants

Twenty-eight participants (mean age 25.93, range 20–36; 12 females) took part in the study, twelve participants per experiment (Exp.1-2-3). Eight participants participated in two experiments. Except from one experimenter, they were naïve to the aim of the experiments and were compensated 10€ per hour. This experiment was approved by the Ethics Committee of the Faculty for Psychology and Pedagogics of the Ludwig-Maximilians Universität München (approval number 13_b_2015) and conducted in accordance with the Declaration of Helsinki. All participants gave written informed consent.

### Apparatus

Participants sat in a quiet and dimly illuminated room. Chin and forehead rests were used to minimize head movements. The experiment was controlled by a PC computer. Gaze position of the dominant eye was recorded using a tower mounted EyeLink 1000 (SR Research Ltd., Ontario, Canada) at a sampling rate of 1000 Hz. The experimental software controlling the display and the response collection as well as eye tracking was implemented in Matlab (The MathWorks, Natick, MA) using the Psychophysics Toolbox (45, 46) and EyeLink Toolbox (47). Stimuli were presented at a viewing distance of 60 cm, on a VIEWPixx LCD monitor (515 by 290 mm, VPixx Technologies Inc., Saint-Bruno, Canada) with a spatial resolution of 1,920 by 1,080 pixels and a vertical refresh rate of 120 Hz. The monitor luminance was linearized with a Minolta CS-100 luminance meter and a custom software (Minolta, Osaka, Japan). Participants’ responses were recorded via a standard keyboard and mouse.

### Procedure

The experiments started with the isoluminance assessment (see below) followed by 8 blocks of the orientation discrimination tasks (Exp. 1, 2 and 3) ran in a single experimental session of about 100 minutes (including breaks).

### Orientation discrimination tasks

Each trial began with participants looking at the fixation point at the screen center (Figure 1A). The fixation point was a “bull’s eye” with a radius of 0.4 degrees of visual angle (dva), composed of superimposed black (~0 cd/m) and white (~57 cd/m^2^) disks. When the participant’s gaze was detected within a 2.0 dva radius of a virtual circle centered on the fixation point, for at least 200 ms, the trial began with a fixation period randomly between 400 and 700 ms. After this period, the presaccadic grating was presented at a distance of 8.0 dva to the left or to the right of the screen center. Participants were instructed to move their eyes as quickly and as accurately as possible towards the grating center. We detected online the saccade onset as the time at which the gaze position left the 2.0 dva radius virtual circle around the fixation point. We next presented the postsaccadic grating (no blank condition, half of the trials) either immediately after the saccade onset detection, or after a 200 ms blank gray screen (blank condition, half of the trials). The postsaccadic grating was presented for 300 ms with a different orientation than the presaccadic grating. Pre- and postsaccadic gratings were green and gray square-wave gratings of 3.2 cycles/dva contained within a circular Gaussian envelope (SD = 3.0 dva). The background was gray and of a different luminance across experiments (Exp. 1: 15 cd/m^2^; Exp. 2: 8 cd/m^2^; Exp. 3: 15 cd/m^2^). The gray stripes of the grating were of the same color as the background. The green stripes of the gratings could either be iso- or anisoluminant relative to the background. The isoluminance of the grating was determined for each participant individually (see *Isoluminance assessment* below). Anisoluminant gratings were having a twofold (~30 cd/m^2^, Exp.1) or fivefold (~40 cd/m^2^, Exp.2) higher luminance as compared to the isoluminant, adjusted value, or being modulated by 6, 12, 25 or 50% as compared to it (Exp. 3). The angle of the presaccadic grating was chosen randomly around the vertical orientation (0°) between ±64°, ±54.5°, ±45°, ±35.5°, ±26° (positive and negative angles refer to clockwise and counterclockwise angles, respectively). The postsaccadic grating was randomly rotated by ±1°, ±4°, ±7°, ±10°, ±13° or ±16° relative to the presaccadic grating angle. Participants reported the change in orientation angle of the postsaccadic grating relative to the presaccadic one. To do so, they used the left or the right direction arrow keyboard buttons, corresponding to clockwise or counterclockwise orientation changes, respectively. Responses were followed by a negative-feedback sound in the case of an incorrect response. In Exp. 1 and 2, presaccadic and postsaccadic gratings were either isoluminant or anisoluminant relative to the background, resulting in four distinct combinations of stimuli across saccades (Figure 2A). In Exp. 3, the pre- and postsaccadic gratings were always of the same luminance in a given trial.

Correct fixation resulted from gaze being maintained within a 2.0° radius virtual circle centered on the fixation target. Correct saccades resulted from saccades landing within a 2.0° radius virtual circle centered on the cue. Trials in which, participants incorrectly fixated, made an incorrect saccade, executed their saccades before than 50 ms or than 350 ms after the presaccadic grating onset or, blinked were repeated. On average 171.53 ± 14.60 trials per participant (11.91 ± 1.01%) were repeated according to these criteria (Exp. 1: 116.50 trials = 11.42 %; Exp. 2: 199.58 trials = 13.86%; Exp. 3: 150.50 trials = 10.45%).

### Isoluminance assessment

Before each experiment, participants performed a block of 40 trials, in which, using the heterochromatic flicker photometry method (48), we evaluated individual isoluminance thresholds of the green stripes of the grating relative to the background. The method consists of presenting the grating with a polarity reversal (±180° phase change) at a frequency of 30 Hz to produce a strong perceived stimulus flicker as long as the luminance of the grating green stripes doesn’t match perceptually with the background luminance. Participants were instructed to minimize the perceived flicker using a computer mouse to adjust the luminance of the green stripes relative to the background. When the flicker was perceived as minimal, they were instructed to press the keyboard space bar. The minimum and the maximum adjustable luminance of the grating green stripes were set between 5 and 25 cd/m² (Exp. 1 and Exp. 3) and between 3 and 13 cd/m² (Exp. 2). The grating was presented in half of the trials at the screen center that participants fixated. On the other half of the trials the grating was shown 8.0 dva to the left or right of the screen center and participants fixated the central bull’s eye. For each participant, a foveal and peripheral threshold of subjective isoluminance was obtained. On average, foveal and peripheral thresholds were 15.47 ± 0.28 cd/m^2^ and 15.83 ± 0.25 cd/m^2^ (Exp. 1), 8.91 ± 0.42 cd/m^2^ and 8.89 ± 0.45 cd/m^2^ (Exp. 2), and, 15.00 ± 0.29 cd/m^2^ and 15.17 ± 0.39 cd/m^2^ (Exp. 3), respectively.

### Data pre-processing

Before proceeding to the analysis of the behavioral results, we scanned the recorded eye-position data offline. Saccades were detected based on their velocity distribution using a moving average over twenty subsequent eye position samples (49). Saccade onset and offset were detected when the velocity exceeded and fell behind the median of the moving average by 3 SDs for at least 20 ms. We included trials where a correct fixation was maintained within a 2.0° radius centered on the fixation target, where a correct saccade started at the fixation point and landed within a 2.0° radius centered on gratings. With such analysis we found that the presaccadic grating was extinguished and replaced by the postsaccadic grating or a blank on average 19.63 ± 0.14 ms after the saccade onset and 29.75 ± 0.75 ms before the saccade offset. In total, we included on average per participants 1388, 1412 and 1229 trials (98.39%, 98.06% and 85.35% of the online selected trials, i.e., 86.29%, 86.12% and 77.33% of all trials played) of the orientation discrimination task of the Experiment 1, 2 and 3, respectively.

### Behavioral analysis

The data for the leftward and rightward saccades were collapsed and percentages of correct responses were calculated for each of the seven different angle changes used across all the presaccadic orientations. Subsequently, cumulative Gaussian functions were fitted to that data using the Palamedes toolbox (50) and the 75% correct discrimination thresholds were determined for each condition and participant. We computed a blanking effect ratio by dividing the orientation threshold observed within trials in which we presented a blank to those observed in trials without a blank and changing this ratio in an improvement percentage, i.e., 100-(x_Blank_ *100 / x_No-blank_). To estimate the overall participant report bias and just noticeable differences (JNDs), we fitted cumulative Gaussian functions to the proportion of clockwise report as a function of the orientation angle change between the pre- and postsaccadic grating. JND angles correspond to the difference between the orientation change needed to obtain 75% and 25% of clockwise reports. Bias angles correspond to the point of subjective equality (PSE), that is the orientation change that would lead to 50% of clockwise reports.

For statistical comparisons 10,000 bootstrap samples were drawn (with replacement) from the original pair of compared values. Then, the difference of these bootstrapped samples was calculated. Finally, two-tailed p-values were derived from the distribution of these differences.

## Author contributions

Conceptualization, design, writing, L.G., H.D. and M.S.; Methodology and analysis, L.G. and M.S.; Resources, H.D. and M.S.; Investigation, L.G.

## Acknowledgements

We thank Jasna Martinovic for her useful comments and, Vanessa Fetzer and Dorontina Ismajli for help with data collecting. This work was supported by grants of the Deutsche Forschungsgemeinschaft (DFG) to HD (DE336/6-1) and MS (SZ343/1) and a Marie Sklodowska-Curie Action Individual Fellowship to MS (704537).

